# Neural correlates of boundary extension during visual imagination

**DOI:** 10.1101/2025.01.31.635941

**Authors:** Timothy Vickery, Banjit Singh, Alyssa Levy, Kallie Sweetman, Zoe Cronin, Helene Intraub

## Abstract

People typically remember seeing a greater expanse of a scene than was visible in a studied close-up (boundary extension, BE). Multivoxel pattern analysis was used to test the neural correlates of BE. Classifiers were trained using a whole-brain searchlight method to discriminate between close-up and wider-angle versions of 16 scenes during repeated perceptual exposures. Earlier, each subject studied either the close or wide version of each scene and then visually imagined it from memory. If a brain region reflects BE, then unlike classification during perception, visual images of close views should sometimes be misclassified as wide (capturing false memory beyond the view), whereas visual images of wide views should be correctly classified. BE-consistent patterns during imagery were found in high-level visual regions, including posterior superior parietal cortex. This pattern did not reflect a brain-wide bias toward better classification of wider-angle views: the pattern reversed (better classification of close views) in the early visual cortex, presenting a novel distinction between early and late visual representations in imagery. We propose that this method reflects active maintenance of boundary-extended scene representations in memory and that it holds promise as a general purpose tool for decoding false memory in the brain.

## Neural correlates of boundary extension during visual imagination

Visual imagery plays multiple roles in cognition; supporting future planning (Bocchi et al., 2011), creativity (Palmierio, Cardi, & Belardinelli, 2011), and memory (Keogh & Pearson, 2011). The current research is focused on image maintenance in working memory (WM), when subjects retrieve a previously studied close-up view of a scene. It has long been known that simple views, studied for a relatively long time (e.g., 15 s each) tend to be misremembered as having shown more of the scene that was shown (boundary extension; Intraub & Richardson, 1989; Hubbard, 2024). This extrapolated content has been observed in drawings of recalled views (Intraub & Richardson, 1989), in responses to test pictures presented in recognition/rating tasks (Intraub, Bender, & Mangels, 1992; Intraub & Richardson, 1989), and in tasks requiring reconstruction of boundary placement (Intraub, Hoffman, Wetherhold, Stoehs, 2006). The goal of the study was to focus on the retrieved image maintained in WM without the presence of a mediating task, such as drawing the image from memory, and without engaging perception, by presenting a test image. We sought to determine if we could decode the neural representation of a boundary-extended visual image in WM using multivoxel pattern analysis (MVPA) to classify neural activity as either matching that associated with the same view during perception or a more wide-angle (more expansive) view of the same scene.

Our study addressed two unknowns about the spatial attributes of the mental image and its neural underpinnings. First, focusing on perception, can a classifier distinguish between a close-up and a slightly wider-angle view of the same scene? Prior research demonstrated that given a small set of natural views, a classifier could discriminate amongst several viewed scenes that differed in category (e.g., the “beach scene” versus a “desert scene”; Johnson & Johnson, 2014). However, in the closer and wider view of, say, a cow in a field, drawing a distinction is more challenging. Here the two views share much of the same visual content, share semantic content, and differ only in terms of expanse (e.g., a bit more of the grassy field). Thus, first, we needed to determine if during perception of closer and wider-angle views, the classifier could distinguish between the two. If so, then the second unknown, which motivated our study, was whether when based on a simple verbal cue (“cow scene”), the neural presentation of the close-up image maintained in WM would match that evoked during perception of the view (the stimulus), or a more wide-angle view that included extrapolated space? To explain our rationale, and the broader implications of our study, we will begin by discussing the brain areas implicated in image maintenance in WM for a variety of stimuli, and then discuss relevant aspects of boundary extension and what is currently known about its neural underpinnings.

Visual imagery induces a BOLD response in occipital, ventral temporal, and parietal visually-responsive regions (e.g., drawings of objects: Ganis, Thompson, & Kosslyn, 2004). Imagined information can be decoded using multi-voxel pattern analysis (MVPA) approaches, which has been demonstrated using a wide variety of stimuli: gratings (Albers, et al., 2013), letters (Stokes, Thompson, Cusack, & Duncan, 2009), categorical (Reddy, et al., 2010), and object stimuli (Ragni, et al., 2020), suggesting common substrates for perception and imagery. Even early visual cortical regions carry information about the contents of imagery (Li, et al., 2023). Visual working memory (VWM) contents, closely related to imagery, are also decodable from even early visual regions (e.g., gratings: Harrison & Tong, 2009).

Such observations have sparked debates about the roles of these regions in these processes, given that VWM and imagery contents are clearly dissociable from perceptual experience. While some accounts posit that frontoparietal regions maintain coarse, non-sensory (e.g., categorical or rule) representations and provide feedback to early visual regions, which maintain specific feature codes used to represent details during VWM and imagery (Sreevnivasan, et al., 2014; D’Esposito & Postle, 2015), other evidence suggests that parietal and frontal regions actually do encode various stimulus-specific properties (Ester, Sprague, & Serences, 2015), and that posterior parietal regions encode qualitatively different properties that are more task-specific and goal-directed than those in early visual cortex (Vaziri-Pashkam & Xu, 2017; Xu, 2018).

Boundary extension provides a new way of exploring this controversy, because boundary extension is a common representational error involving spatial expanse of a view; for object-based close-ups it is highly reliable, evident following relatively long study times (15 s; e.g., Intraub et al., 1992; Munger & Multhaup, 2016) and brief study times (250 ms; Bainbridge & Baker, 2020; Intraub & Dickinson, 2008). It has also been observed in memory for scene-regions in real space following either visual or haptic perception (Intraub, 2004; Intraub et al., 2015; Mullally et al., 2012). Pictures need not encompass a semantically plausible view to elicit boundary extension (e.g., snow skier in a parka on a sunny beach; Mamus & Boduroglu, 2018), an important characteristic of boundary extension is that the picture is interpreted as a view of a continuous scene.

A striking example that is particularly relevant to the current research, involved explicit imagination instructions (Intraub, Gottesman, & Bills, 1998). When line-drawings of *only* the main object were traced from close-up or more wide-angle views of a scene, memory errors did not reflect boundary extension. Although subjects’ task was to remember the size and placement of the object in the display, large objects (e.g., a traffic cone traced from the close-up view) were remembered as having been a little smaller, whereas small objects (e.g., the traffic cone traced from a very wide-angle view) were remembered as a little larger (consistent with normalization in memory). However, when the same stimuli were presented to other subjects, who in addition to the same instructions, were asked to imagine a background around the object (e.g., for the cone: the background was asphalt, with a shadow to the left), “projecting it” onto the display to “aid memory”, the error pattern changed, reflecting the common pattern associated with boundary extension – the larger objects were remembered as having been much smaller, and the smaller objects elicited no directional error. In fact, mean boundary ratings in the imagine-background condition were indistinguishable from those elicited by the same line-drawn objects with the backgrounds sketched into the display. Thus, in the critical conditions, given the same line-drawn objects, the mental representation differed depending on whether or not the visual stimulus had been interpreted (and imagined) as a view of a scene. In the current study, we reasoned that the retrieved visual image (including boundary extension) of a close-up photograph would differ from the perceived image in a predictable way – reflecting a more expansive view of the scene than had been shown, and we sought to determine if this could be detected in the neural representation.

Prior research addressing different questions have been successful in detecting areas of the brain associated with boundary extension. Park et al. (2007) used a repetition attenuation paradigm in which close or wider views, either repeated by presenting the identical view or repeated by presenting the alternative view. When different views were shown, both RSC and PPA demonstrated the response asymmetry commonly observed in behavioral studies. Although responding to the same comparison (close vs wide), when the close view was presented first, presentation of the wider view later, was met with repetition attenuation; whereas when the wider view was presented first, there was no repetition attenuation in response to the close-up. In other words, both brain areas had “misjudged” the close-wide views as being the same view, whereas they correctly responded to the wide-close views as different views. When the two views were identical, the response in RSC was consistent with boundary extension, no repetition attenuation was observed, whereas in PPA attenuation was observed. Thus, RSC, an area that appears to be associated with thinking about a view within in greater geographical context (Epstein & Baker, 2019) appeared to be responsive to boundary extension in memory, and PPA partially responsive (clearly so, which respect to the mismatch conditions). This was thought to be due to the PPA being responsive to layout but also identity.

In other research Chadwick et al. (2013) examined activity in response to the initial presentation of a scene, and using univariate analyses compared trials on which BE was subsequently observed in behavior vs. when it was not. They found evidence that the hippocampus showed elevated activity associated with subsequent earmarks of BE, consistent with neuropsychological studies of subjects with brain damage (De Luca et al., 2018; Maguire et al., 2016; Mullally et al., 2012 ). Additionally, adaptation effects were observed in early visual areas of occipital cortex, which were stronger for trials on which BE was not observed than those in which it was observed. This implies a role of the early visual cortex in representing a boundary-extended scene.

Both Park et al. and Chadwick et al. examined BE correlates using univariate approaches. Additionally, both relied on perception – either detecting the neural response during stimulus presentation (Chadwick et al.), or the adaptation effect in response to presentation of the test picture (Chadwick et al. and Park et al.). Neither queried the properties of the retrieved image maintained in WM, which is the aim here, enabled by the use of MVPA.

In this study, we sought neural evidence of BE when subjects retrieved and actively maintained an image of a studied scene in WM. The method is illustrated in Figure 1A. In Stage 1, subjects studied either the close or wider-angle version of a scene, then a short time later each view was cued with a word describing the main object, and subjects were instructed to “project” an image of that view into a rectangle in the same location that the original stimulus had been presented (to control the scale of their image); fMRI data was collected during this imagery phase. In Stage 2, the stimuli were presented again and subjects rated them as the closer or farther (4-point scale) to obtain a behavioral index of memory for the view’s expanse in a common recognition memory task. Later, in Stage 3, to train a classifier, they repeatedly viewed the close and wide versions of the same scenes and fMRI data was collected during this perceptual phase. In our key analysis, a classifier was trained on Stage 3, and its performance in determining whether the imagined view was close or wide was determined on Stage 1 Imagery trials. In a brain region that reflects a boundary-extended visual image during retrieval and maintenance in WM, we would be expect the classifier to err on trials in which subjects had studied the close-up view, as compared with the wider-angle view. Regions that do not reflect an extended image would be expected to do as well at decoding memory for close and wide views during image maintenance. Other regions, reflecting qualitatively different distortions, might make more errors in the recall of wider-studied views.

**Figure 1.**
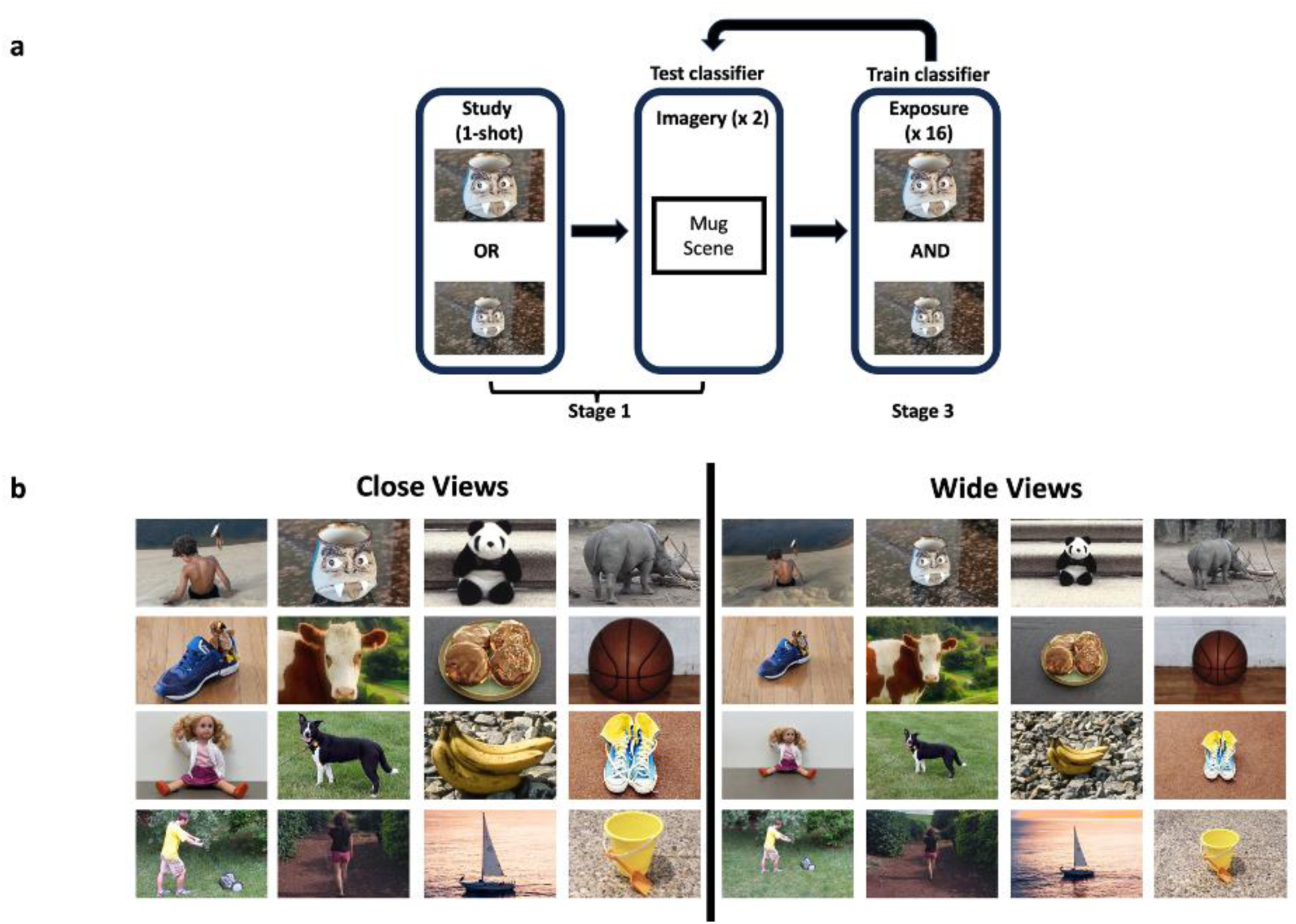
A) A schematic of the basic design logic. Images (half presented in close views and half in wide views) were studied and then imagined. Later, close and wide views were presented. A pattern classifier was trained to discriminate close vs. wide presentations, then tested on its performance during imagery trials. B) The 16 scenes, in close and wide views, used in this study.

In contrast to prior efforts to examine the neural correlates of BE, this method interrogates the contents of imagined memories, rather than the brains’ response to either the initial image presentation or a second presentation of the image. Thus, it more directly engages the representation of the recalled scene.

## Materials and Methods

### Subjects

Subjects were 21 (11 female, 10 male) members of the community surrounding the University of Delaware, ages 18 to 43. Subjects were right-handed and had normal or corrected-to-normal vision. The study was conducted under the approval of the University of Delaware Institutional Review Board. All practices conformed to the Declaration of Helsinki, with subjects providing informed written consent to the experimental protocol. One subject’s data was excluded from analysis due to high signal dropout in MRI scans (visible distortions and signal dropout in raw images).

### MRI procedures

MRI data were collected using a Siemens 3T Magnetom Prisma scanner with a 64-channel head coil.

#### Scanning protocols

##### Structural scan

T1-MPRAGE scans were collected following standard localizer scans. Images were acquired with 0.7 mm on edge isotropic voxels (TE = 46.4 ms, TR = 2.08 s, FA = 9°).

##### Functional protocol

Functional scans were whole-brain T2*-weighted images collected using a multi-band (MB factor = 8) echo-planar imaging sequence consisting of 72 slices with an oblique axial orientation acquired with a resolution of 2 mm by 2 mm by 2 mm (sequence parameters: TR=1 s, TE = 40 ms, FA = 52°). Initial images (prior to trigger from scanner) in each scan were automatically discarded.

### Stimuli

A total of 16 unique scenes were employed; a close-up and a wide-angle version of each were created. To achieve our goals, the optimal wide view would be sized to mimic boundary extension for the close view. Boundary extension varies across subjects and pictures, and thus of course, cannot be determined in advance. We decided to base sizing of the wide views on subjects’ drawings from memory in Intraub and Bodamer (1993) because this is a recall task, and like our stimuli, these were object-based views. We were able to create a wide view based the reflected the mean change in area of the main object in the boundary extended drawing of that picture for five of our stimuli. For the rest (new object-based scenes), in one case, the view was similar to a stimulus in Intraub and Bodamer, so we sized it in relation to those drawings, and for the rest, we sized the image to match the mean across the drawings selected. On average, in 16 scenes, the area of the main object in the wide view was reduced to .38 (SD = .04) of its area in the close view; across scenes the area was reduction of the main object ranged from .26 to .46.

### Task design

Behavioral tasks began following the T1 scan. Tasks completed inside the scanner were divided into three stages. Prior to scanning, each subject completed a two-image version of Stage 1, to ensure that they understood the primary task. The two images used in practice were not used for scanning runs. While scanned, subjects held a 4-button response box in their right hand for making responses.

#### Stage 1: Study and Imagine

In the first stage (Figure 2A), subjects studied and then imagined 4 unique scenes in each of 4 runs while scanned (duration: 244 s each run), for a total of 16 unique scene .8 of these were presented using the close version and 8 using the wide-version. Each run consisted of 2 close and 2 wide views, presented either close-wide-close-wide or wide-close-wide-close, with that order alternating run-to-run; the starting order was counterbalanced across subjects. The scenes were divided into two sets; one set was presented as close to half the subjects and the other set was presented as close to the other half. The order of presentation of images was randomized within the above constraints.

**Figure 2.**
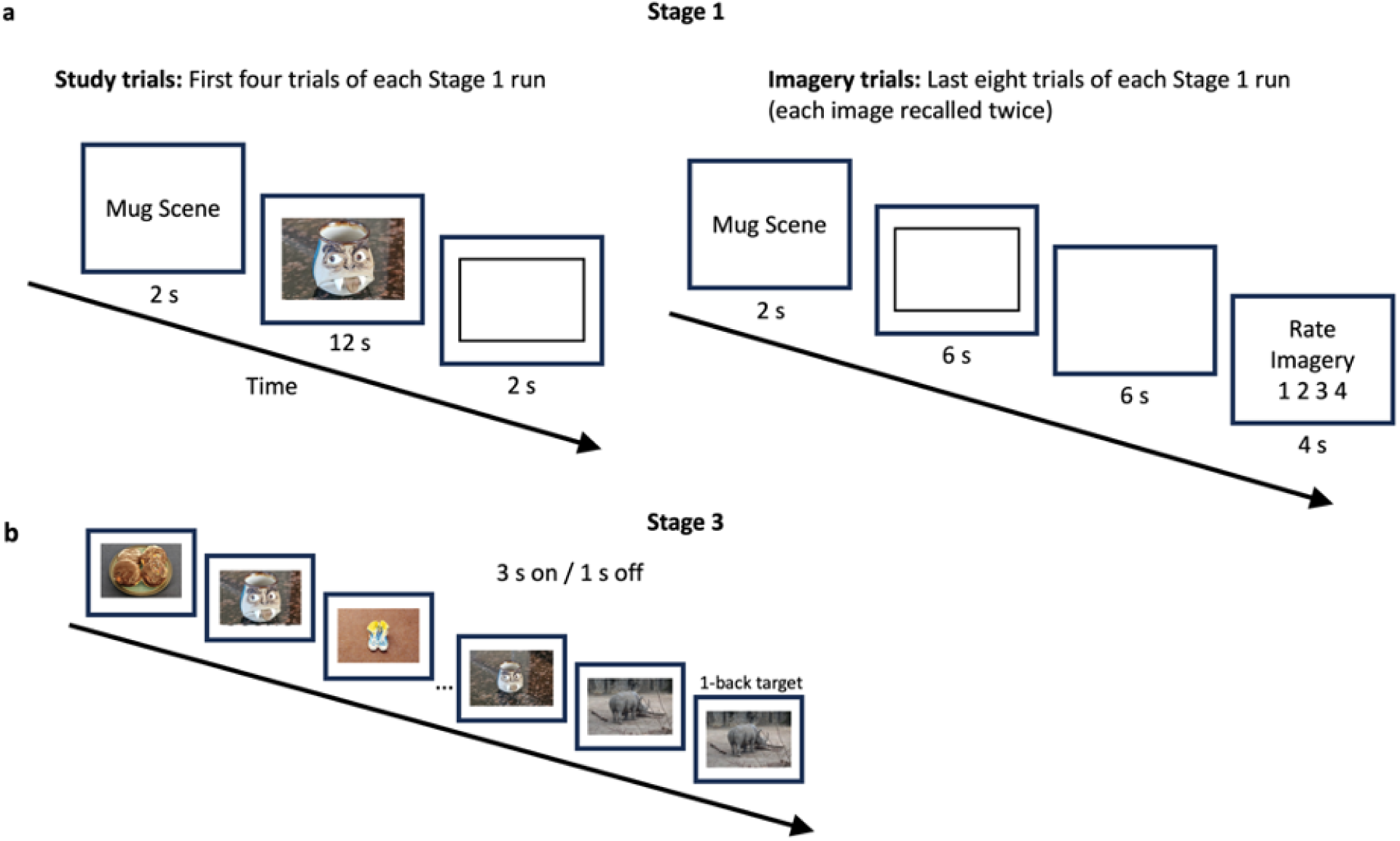
A) Stage 1 design. Each image was studied once for 12 s. In the same run, the image was later recalled and imagined, twice, cued by its label. B) Stage 3 design. All close and wide views presented multiple times (3 s on, 1 s off, except for 3 subjects – see text).

Within each run, after 4 s delay after scanning was triggered, a label (e.g., “MUG SCENE”) appeared for 1.5 s followed by a 0.5 s break, followed by 12 s appearance of the associated view. After a 2 s break, the next trial’s label appeared.

After the offset of the 4th view and a 2 s blank, a message was presented for 6s that read, “Short break…get ready to imagine! Project the image as vividly as possible into the rectangle when cued with the label!” A 6 s blank followed before the onset of the first label cued which scene to imagine (e.g., “MUG SCENE”). After a 1.5 s phrase cue and a 0.5 s break, a black frame (the same size as the originally studied image) appeared for 6 s and subjects projected their image in this space. A 6 s break followed and then, for 4.0 s, an imagery rating cue was presented. Subjects were instructed to rate their imagery on a 4-point scale (very weak, weak, strong, very strong) using a four-button box following every imagery trial. Following the rating screen a 2 s blank preceded the onset of the next label.

In the same order as the study phase, each memory was cued for imagery once. Then, each image was cued again in the same sequence. Thus, each studied image was associated with 2 recall and imagine events. After the final rating, a 6 s blank period completed the run.

#### Stage 2: Scene memory test

To obtain an index of boundary extension, we next presented the studied views one at a time and instructed subjects to rate whether the view appeared much closer, closer, farther, or much farther than originally studied. Trials were unspeeded and no MRI data were acquired during this stage.

#### Stage 3: Close and Wide Views (Classifier Training Trials)

In this stage, both the close and wide views of each scene were each shown twice per run There were 8 runs. Each of the 32 views was presented for 3 s followed by a 1 s blank in a randomized order with the constraint that the same scene could not repeat, even in the alternative view, back-to-back. Two 1-back trials were inserted per run (these were removed for classification analysis), and the subject’s task was to respond with a button press when this repetition occurred. Null trials (2) were randomly inserted into each run, with no image presented. A 4 s blank preceded the run and a 10 s blank terminated the scan. Each run lasted 302 s.

For the first 3 subjects scanned (one of whom was excluded due to image distortion, see above), the timing of this stage differed. Images were presented for 2 s and followed by a variable delay of between 2 and 5 seconds (mean inter-trial interval was 3.5 s). There were no null trials and the scan lasted 382 s. Otherwise, it was the same as the revised protocol. These subjects completed 6 scans of this protocol. The protocol was altered for the following reasons: (1) the timing characteristics caused subjects boredom and fatigue; (2) random jittering is not useful in extracting single-trial estimates; and (3) an additional two scans were made possible with the revised protocol. The revised protocol was better tolerated, and we deemed it to likely provide better single-trial estimates. Analysis of these trials was adapted to timing and duration accordingly.

#### Functional localizer

A separate functional localizer run presented short 3 s movie clips of scenes, faces, bodies, moving inanimate objects, and scrambled moving objects, crossed with color or grayscale. Blocks of 6 clips (lasting 18 s) of each class were presented in an AB..BA order, with a blank 16 s interval at the beginning and end. This scan was typically conducted in the middle of Stage 3 runs, to disrupt monotony, and lasted 450 s.

### Data analysis

Primary analysis involved the training of support-vector machine classifiers to discriminate close and wide views of each individual scene (from Stage 3 data), separately, then testing the classifier on corresponding Stage 1 imagery trials. Performance was averaged separately for close and wide.

#### Preprocessing

Preprocessing was conducted using AFNI (Cox, 1996; Cox & Hyde, 1997). All volumes were motion-corrected, smoothed with a 4-mm FWHM kernel, aligned to the structural images, and corrected to MNI152 2mm standard space (via structural registration to standard). Voxel-wise, activity was scaled within run.

#### Single-trial estimation

For classification, individual trial responses were extracted using GLMSingle (Prince et al., 2022) with preprocessed images as inputs. This was done separately for Stage-1 imagery trials and for Stage-3 perception trials, employing a 3 s and 6 s trial duration, respectively. Note that temporal filtering was not conducted at preprocessing, as GLMSingle employs low-order polynomial regressors to accomplish the same ends.

#### Classification

Classification was conducted using a searchlight procedure implemented in CosmoMVPA (Oosterhof et al., 2016), and on an ROI-by-ROI basis using individual subject ROIs defined by analysis of the functional localizer. For the searchlight procedure, we requested 179-voxel searchlight regions, corresponding to a 7-mm radius sphere with our image resolution. The classifier we employed was Linear Discriminant Analysis (LDA).

A leave-one-run-out cross-validation procedure was used to assess the viability of decoding close vs. wide views of scenes from Stage 3 perception data. This established the extent of regions within which the classifier could decode close vs. wide view during perception of the view, and therefore, established the regions within which we could reasonably expect to observe decoding during imagery. Wide vs. close classifiers were trained and tested for each scene, separately. Accuracy maps were formed for each scene and then averaged within each subject. Across subjects, the maps were contrasted to chance with a one-sample t-test at each voxel. Searchlight statistical maps were combined across subjects and cluster-corrected using threshold-free cluster enhancement (Smtih & Nichols, 2009) as implemented in CosmoMVPA using 10000 permutations. To examine whether there was a bias due to close vs. wide (Stage 1) study assignment, we also split accuracy maps according to study condition, and contrasted those with a paired t-test. To examine whether there was a bias based upon the decoded image (i.e., whether it was easier to classify an image as wide or close, overall), we conducted the same split based upon test item (wide or close).

For the imagined view classification, our primary target analysis (train on Stage 3, test on Stage 1 imagery), results were partitioned according to whether the item was studied as a close or wide view. The classifier, trained on all Stage 3 runs to discriminate wide and close views during perception, was applied to single-trial estimates extracted from Stage 1 imagination trials. Accuracy was averaged for scenes studied wide or close, separately. Wide and close maps were compared to chance performance (50%), separately, and contrasted against one another.

### Functional localizer

This scan was processed using AFNI, with the same preprocessing protocols but also including a typical regression design in which blocks were modeled. Contrasts of interest were scenes > other categories, and in post-hoc analysis, face > other categories. ROIs were localized with two methods. (1) A peak voxel in the approximate region of PPA or RSC in each hemisphere was isolated on a per-subject basis, and a 8-mm radius sphere was taken, and (2) using the group-constrained, subject-specific approach suggested by Fedorenko et al. (2010). The resulting regions were employed in classification analysis. We only pursued Stage 3 to Stage 1 transfer analysis if Stage 3 leave-on-out analysis performed better than chance.

## Results

### Behavior

Ratings from Stage 2 were coded as -2 (much closer), -1 (closer), +1 (farther), and +2 (much farther). Our response box had only four buttons, and therefore no 0 (same) option was included. Results split by the original studied view are shown in Figure 3. Close views were rated as closer than originally studied (M = -0.68, SD = 0.60) and were significantly different from a mean of 0, t(20) = -5.17, p < .001, d = -1.13. Wide views (M = -0.18, SD = 0.66) were not significantly rated differently from 0, t(20) = -1.28, p = .21, d = -0.28. The close view ratings were significantly lower than wide view ratings, as well, t(20) = -4.64, p < .001, d = 1.01.

**Figure 3.**
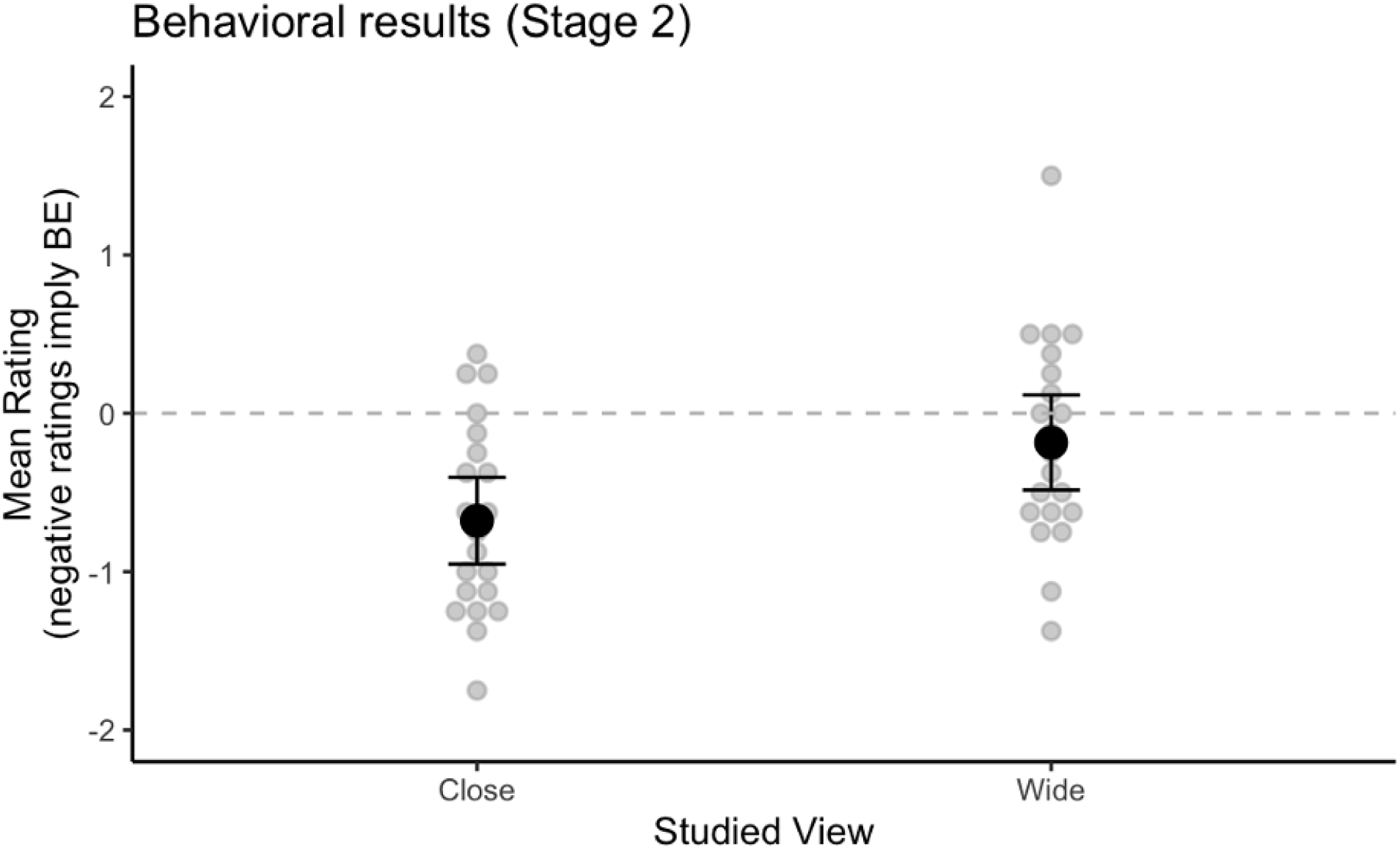
Individual (gray dots) and mean (black dots) performance in Stage 2, split according to whether the scene was originally studied in the close or the wide view.

These results support the presence of significant BE in close-studied images, which showed both significant BE, and were significantly greater than wide-studied views. No significant BE was observed for wide-studied images.

### Decoding perception of scene expanse

Our first major approach to analysis of the fMRI data was to consider the ability of a classifier to distinguish between close and wide views of the same scene. We reasoned that testing a classifier trained on perception of scenes in its ability to discriminate close and wide view memories (train on Stage 3, test on Stage 1 imagery) would not be sensible if the classifier could not discriminate between close and wide views during perception (Stage 3 alone).

The searchlight procedure was performed on a per-scene basis, and performance was aggregated across scenes by averaging accuracy. The resulting maps were compared to chance (p < .05, one-tailed, TFCE-corrected) to locate regions that performed at above-chance levels. The results in Figure 4A show that most of the occipital cortex, regions of posterior and superior parietal cortex, and regions of posterior inferotemporal cortex did support such classification.

**Figure 4.**
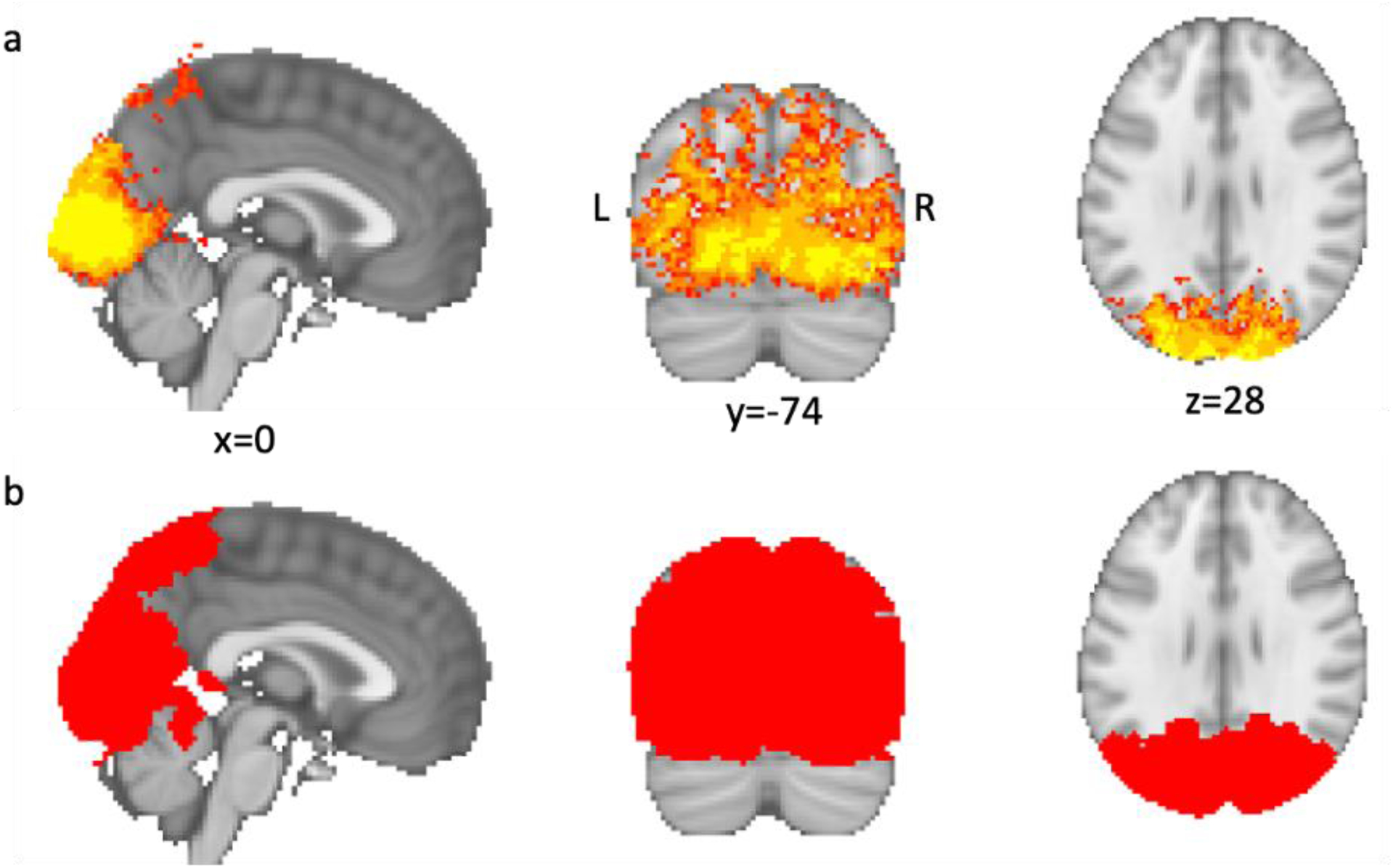
a) Regions highlighted were significantly better than chance (p < .05, one-tailed, TFCE-corrected) at decoding close vs. wide views during perception (Stage 3) in a leave-one-run-out procedure. b) the mask in (a) dilated twice was used as a mask for the searchlight analysis that tested transfer from Stage 3 (perception) to Stage 1 (imagery).

**Figure 5.**
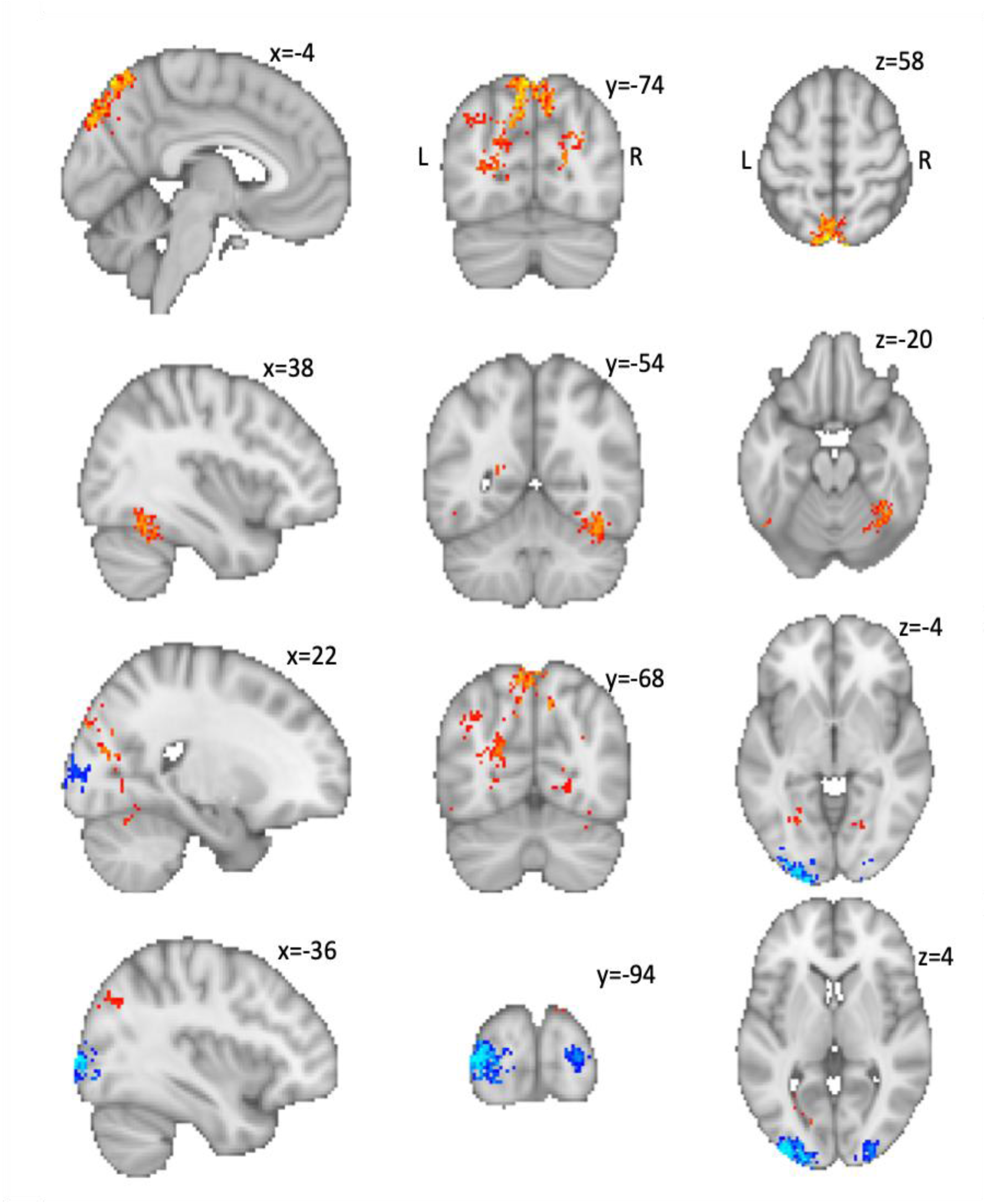
Results of contrasting performance at decoding wide-studied imagery as wide vs. decoding close-studied imagery as close (P < .05, two-tailed, TFCE-corrected). Warm regions showed better performance for wide-studied imagery than close-studied imagery, consistent with BE. Cool regions showed the opposite pattern.

Figure 4B shows a twice-dilated version of a mask formed from the results of the above analysis, intersected with the MNI152 brain mask. This mask was used to constrain the searchlight for our primarily imagery-stage analysis, below.

### Testing for bias in perception decoding

The central test, decoding imagery, depends upon a comparison that could be ambiguous if the classifier at perception shows a bias based on either the manner in which the scene was initially studied (close or wide) or whether the target of classification itself is a close or wide view. Two control analyses were conducted to examine these possibilities.

First, we split perception-stage decoding results into scenes that were studied close and those studied wide and performed a paired t-test at each searchlight center between accuracy for each of these cases. No cluster survived correction (p < .05, two-tailed, TFCE-corrected). This indicates that the initial encounter with the image, during encoding, did not affect the classifier’s ability to decode close vs. wide in perception.

Second, we split the perception-stage decoding results based on the view decoded (close or wide). The goal of this analysis was to locate any regions in which there was a significant bias, for example, to decode wide views and close views as wide. As in the first control analysis, we found no clusters (p < .05, two-tailed, TFCE-corrected) that reflected any such bias.

### Decoding imagery of images studied in wide versus close views

In our primary target analysis, we trained the classifier to discriminate close and wide views (Stage 3), and tested its performance on imagery trials (Stage 1), split by whether the original image recalled was studied in a wide view or a close view. We reasoned that a region with representations that reflect BE should show better performance for wide-studied images than close-studied images.

Within the mask region established by the Stage 3 LOO analysis, we found several regions that were consistent with BE. Four major clusters were found (Figure 4 and Table 1). The largest cluster had peak differences in bilateral superior parietal cortex, and extended into dorsal occipital regions. The next two were located in right fusiform and left inferotemporal cortex. The final cluster was located in a right lingual region.

**Table 1.**
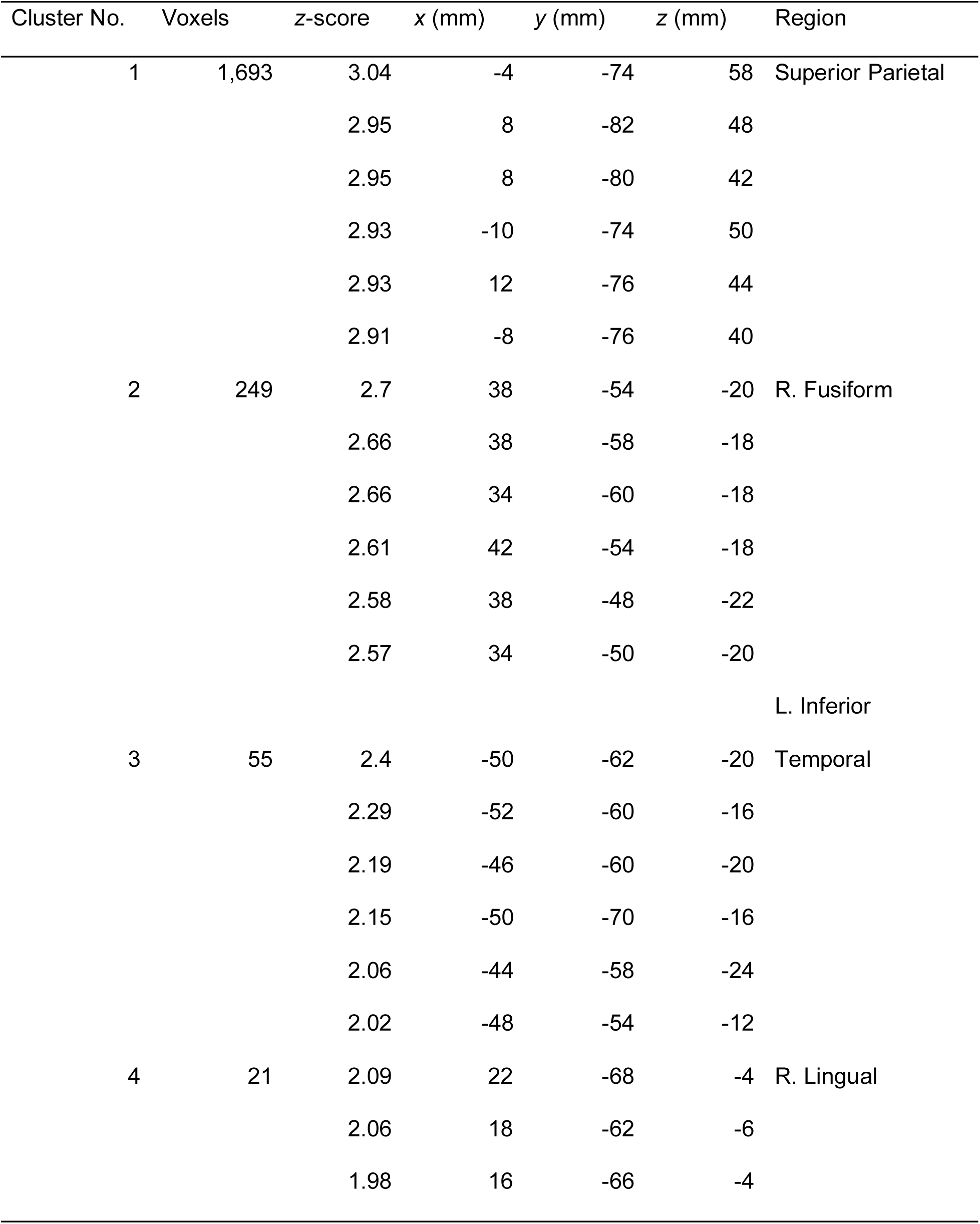
Four clusters that showed significantly greater performance for wide-studied imagery than close-studied imagery.

**Table 2.** Two clusters that showed significantly greater performance for close-studied imagery than wide-studied imagery.

| Cluster No. | Voxels | z-score | x (mm) | y (mm) | z (mm) | Region |
| --- | --- | --- | --- | --- | --- | --- |
| 5 | 701 | 3.54 | -32 | -94 | 10 | L. Occipital |
|  |  | 3.54 | -36 | -94 | 4 |  |
|  |  | 3.54 | -32 | -94 | 6 |  |
|  |  | 3.43 | -26 | -92 | 8 |  |
|  |  | 3.43 | -20 | -102 | -6 |  |
|  |  | 3.43 | -26 | -100 | 6 |  |
| 6 | 199 | 2.91 | 28 | -94 | 4 | R. Occipital |
|  |  | 2.75 | 28 | -100 | 6 |  |
|  |  | 2.69 | 26 | -94 | 0 |  |
|  |  | 2.64 | 28 | -90 | 0 |  |
|  |  | 2.56 | 24 | -100 | 4 |  |
|  |  | 2.46 | 20 | -98 | 8 |  |

The opposite pattern was also observed. Some early visual regions showed better performance classifying close-studied imagery as close than wide-studied imagery as wide. The two clusters spanned regions near left V1, V2, and V3 in a probabilistic map (we did not have retinotopy data for these subjects), and near right V2 and V3.

No region showed significantly worse-than-chance activity (p < .05, TFCE corrected) for either close or wide view memories. The wide-preferring regions overlapped with regions that were above-chance at decoding wide recall as wide, and the close-preferring regions overlapped with regions that were above-chance at decoding close recall as close.

### Regions of Interest (ROI) Analysis

We isolated scene-selective parahippocampal place area (PPA) and retrospenial complex (RSC) in individual subjects, using the localizer task data. We found the peak voxel activity for a scene > other categories contrast within each region and created a 10-mm radius sphere around this voxel (515 voxels), separately for each hemisphere. We then conducted the same MVPA analyses within each of these regions.

Neither of the scene-selective regions showed significant ability (p < .05, uncorrected) to classify views of a scene as close vs. wide. For this reason, we did not conduct the primary analysis of transfer to the imagery task. The same result was obtained when we conducted a group-contrained subject-specific (GSS) analysis (Fedorenco, et al, 2010).

The searchlight data showed clusters in fusiform cortex that showed signatures of boundary extension. Therefore, we conducted a post-hoc analysis in which we isolated the fusiform face area (FFA) from a face > other classes contrast, and created 10-mm ROIs in each hemisphere.

The left FFA was just above chance at decoding close vs. wide views in perception (accuracy M = .512, SD = .024, p = .04 uncorrected vs. chance). The right FFA was also above chance (M = .521, SD = .027, p = .002).

We therefore examined the transfer from perception to imagery in these regions, comparing performance at classifying wide-studied imagery with classifying close-studied imagery. Left FFA did not significantly differ in its performance between these two classes (p = .18). However, right FFA did show a difference that was consistent with BE. Performance for close-studied imagery (M = .431, SD = .143) was worse than chance, meaning that close-studied imagery was misclassified as wide (p = .045), whereas wide-studied imagery was not significantly different from chance (M = .534, SD = .155, p = .33). The difference between the two was marginally significant (p = .045), implying that right FFA representations during imagery showed a BE-consistent pattern, albeit one that differed a bit from our central searchlight findings. That is, in right FFA, close-studied imagery was decoded as wide-studied, above chance levels, whereas wide-studied imagery was at chance. In contrast, our searchlight revealed regions where wide-studied imagery was above-chance, and close-studied imagery was at chance.

## Discussion

Our results suggest that MVPA can be used to decode boundary-extended representations that were retrieved and maintained in working memory. In our study, subjects studied close or wide views of scenes, imagined them from memory, and then later viewed repeated instances of both the close and wide versions of the same studied scenes. Subjects’ boundary ratings indicated the presence of boundary extension for close-up scenes, while ratings of wide-angle scenes did not, consistent with many prior studies using object-based close-ups such as these, which show less BE or no directional error for wide-angle views (e.g., Intraub et al., 1992; see Intraub 2002 and Hubbard et al’s., 2010, review). A classifier trained on perception of close vs. wide views in the final stage of the experiment was applied to the imagery stage. In regions such as anterior occipital cortex, posterior and superior parietal regions, and inferior temporal regions, classifiers made more errors when decoding imagery of close-studied scenes (confusing them more often with wide-viewed scenes) than they did with wide-studied scenes (which were more often decoded as wide-viewed than close-viewed). This pattern of activity putatively reflects the gain of expanse that occurs in memory representations.

In contrast to prior neuroimaging research on boundary extension, which relied on signals at the time of initial study or re-presentation of the varied images, the present study sought to interrogate recalled representations and recover aspects of their distorted representations in memory. As such, this study is the first example of decoding false memories of which we are aware. Unlike previous studies of boundary extension, our goal was to develop a tool for the study of the contents of boundary-extended representations, rather than to test the roles of specific regions in the formation of boundary-extended memories.

Another contribution of the current work is to demonstrate the basic decodability of close and wide views of a single scene. Notably, close and wide views are quite similar in terms of image properties, and thus the viability of a classifier to discriminate between the two based on a small amount of training data was not taken for granted. Indeed, prior work has focused on much broader distinctions, such as categorical differences (e.g., Reddy et al., 2010; Johnson & Johnson, 2014). However, our classifiers accomplished this task in many regions spanning most of the occipital cortex, a large expanse of posterior parietal cortex, and some posterior inferior temporal regions. It did so with no evident bias, as success in decoding close and wide views was not significantly different. This sensitivity to spatial expanse was critical to our effort in decoding mental imagery and may represent a useful tool for investigating other questions.

Ideally, boundary extension would be reflected by reliable worse-than-chance confusion of close-studied imaged scenes with wide views (i.e., classification of close views as wide). While we found that the classifier did poorly in the regions highlighted above, exclusively for close-studied scenes (often confusing them with wide views), we did not find worse-than-chance error rates. One possibility is that the wide views employed in this study were more expansive than the average boundary-extended representations of close views. We chose the wide views based on prior research with images of the same scenes, but those studies employed a longer retention interval and probed memories with drawings. However, the observed pattern was still robust and captures a basic confusion that is *exclusive* to close-studied scenes in these regions.

This issue points to important future directions. First, we plan to calibrate wide views according to a procedure that more carefully matches the experimental context used in our MRI study. Second, future work can more carefully assess individuals’ boundary extension on a per-scene basis, filtering trials with less exaggerated boundary extension and/or correlating neural signals of boundary extension with individual responses.

A limitation of the current study is that we are limited to regions that can decode the rather subtle differences between wide and close views in *perception*. The interpretation of the classifier, when applied to imagery, is unclear in the absence of such an ability. One result of this is that we were limited in interpretation of regions of interest such as RSC and PPA, which had poor classification accuracy for perceptual differences between close and wide. However, the primary utility of this method is to interrogate the contents of memory rather than test hypotheses about the roles of different regions. Therefore, it was critical to focus on sites that showed the sought-after pattern of confusion errors, rather than on specific regions that might be important to the formation of such images. Nevertheless, the regions in which we found the sought-after pattern of confusion errors were sensible in the context of previous research on imagery and working memory. In particular, posterior parietal and anterior/superior occipital regions are previously implicated in the recall and maintenance of imagery and working memory representations (e.g., Ragni, et al., 2020; Albers, et al., 2013).

Interestingly, the tendency to confuse close-studied scenes with wide-viewed image responses was not ubiquitous, even in regions that could successfully discriminate between close vs. wide viewed scenes. In particular, early occipital regions showed the opposite pattern, with classifiers performing better at decoding imagery of close-studied scenes as close views than they decoded wide-studied scenes as wide views. One possible reason for this observation is that it might reflect the importance of focal information in these early visual regions. In particular, early visual regions tend to over-represent the foveal information spatially, moreso than many higher-level visual regions (Arcaro et al., 2009; Wandell et al., 2007). Since object information in these images was central, and most likely foveated, the classifier trained to discriminate close vs. wide views might simply do better for close-studied scenes because most of the important information in these regions is more discriminable in foveal regions for close views.

The distinction between early and late visual regions may also be informative to debates about the roles of early vs. late regions in working memory and imagery. It is well recognized, for example, that early regions re-represent information held in working memory, as well as late regions that are thought to be critical for WM and imagery. The difference in patterns of errors may point to differences in the roles of the information represented in these regions, with information available for explicit recall (and thus possessing boundary-extended contents, in this experiment) represented in later regions.

While our observations of early visual cortex should be the focus of future research, it was fortuitous in the sense that it eliminated one counter-argument for the pattern observed in later visual regions. Specifically, it was *not* the case that the classifier simply did more poorly, everywhere, in decoding imagery of close-studied scenes. This bolsters the argument that the observation was indeed due to a change in representation from study to imagined memory.

In conclusion, our method and findings represent a novel approach to the study of false memories, specifically boundary-extended memories of visual scenes. Along with drawing and other behavioral indices of the gain of expanse observed after studying close views of scenes, this method can be used to probe the contents of remembered scenes. A key advantage to this new method is that, unlike these behavioral indices, it requires no new presentation of the scene (behavioral judgments of re-presented scenes, such as used in our Stage 2), nor does it require explicit interaction with the memory (as in drawing from memory, in which remembered representations interact with the action system to produce the drawing). The key data in our novel method arises only from the effort to remember the viewed scene based upon a simple verbal label, requiring no other action from the observer and avoiding presenting the studied scene again. Thus, this research opens doors to novel questions about boundary-extended representations

## Acknowledgements

This work was supported by NIH NIGMS 5P30GM145765-03.

## References

Albers, A. M., Kok, P., Toni, I., Dijkerman, H. C., & de Lange, F. P. (2013). Shared representations for working memory and mental imagery in early visual cortex. Current biology : CB, 23(15), 1427–1431. 10.1016/j.cub.2013.05.065

Arcaro, M. J., McMains, S. A., Singer, B. D., & Kastner, S. (2009). Retinotopic organization of human ventral visual cortex. The Journal of neuroscience : the official journal of the Society for Neuroscience, 29(34), 10638–10652. 10.1523/JNEUROSCI.2807-09.2009

Bainbridge, W. A., & Baker, C. I. (2020). Boundaries extend and contract in scene memory depending on image properties. Current Biology, 30(3), 537–543. 10.1016/j.cub.2019.12.004

Bocchi, A., Carrieri, M., Lancia, S., Quaresima, V., & Piccardi, L. (2017). The key of the maze: The role of mental imagery and cognitive flexibility in navigational planning. Neuroscience Letters, 651, 146–150. 10.1016/j.neulet.2017.05.009

Chadwick, M. J., Mullally, S. L., & Maguire, E. A. (2013). The hippocampus extrapolates beyond the view in scenes: an fMRI study of boundary extension. Cortex; a journal devoted to the study of the nervous system and behavior, 49(8), 2067–2079. 10.1016/j.cortex.2012.11.010

Cox R. W. (1996). AFNI: software for analysis and visualization of functional magnetic resonance neuroimages. Computers and biomedical research, an international journal, 29(3), 162–173. 10.1006/cbmr.1996.0014

Cox, R. W., & Hyde, J. S. (1997). Software tools for analysis and visualization of fMRI data. NMR in biomedicine, 10(4-5), 171–178.

De Luca, F., McCormick, C., Mullally, S. L., Intraub, H., Maguire, E. A., & Ciaramelli, E. (2018). Boundary extension is attenuated in patients with ventromedial prefrontal cortex damage. Cortex, 108, 1–12. 10.1016/j.cortex.2018.07.002

D’Esposito, M., & Postle, B. R. (2015). The cognitive neuroscience of working memory. Annual review of psychology, 66, 115–142. 10.1146/annurev-psych-010814-015031

Epstein, R. A., & Baker, C. I. (2019). Scene Perception in the Human Brain. Annual review of vision science, 5, 373–397. 10.1146/annurev-vision-091718-014809

Ester, E. F., Sprague, T. C., & Serences, J. T. (2015). Parietal and Frontal Cortex Encode Stimulus-Specific Mnemonic Representations during Visual Working Memory. Neuron, 87(4), 893–905. 10.1016/j.neuron.2015.07.013

Fedorenko, E., Hsieh, P. J., Nieto-Castañón, A., Whitfield-Gabrieli, S., & Kanwisher, N. (2010). New method for fMRI investigations of language: defining ROIs functionally in individual subjects. Journal of neurophysiology, 104(2), 1177–1194. 10.1152/jn.00032.2010

Ganis, G., Thompson, W. L., & Kosslyn, S. M. (2004). Brain areas underlying visual mental imagery and visual perception: an fMRI study. Brain research. Cognitive brain research, 20(2), 226–241. 10.1016/j.cogbrainres.2004.02.012

Harrison, S. A., & Tong, F. (2009). Decoding reveals the contents of visual working memory in early visual areas. Nature, 458(7238), 632–635. 10.1038/nature07832

Hubbard, T. L., Hutchison, J. L., & Courtney, J. R. (2010). Boundary extension: Findings and theories. Quarterly Journal of Experimental Psychology, 63(8), 1467–1494. 10.1080/17470210903511236

Hubbard T. L. (2024). Setting the scene for boundary extension: Methods, findings, connections, and theories. Psychonomic bulletin & review, 10.3758/s13423-024-02545-w

Intraub, H. (2002). Visual scene perception. In Nadel, L. (Ed.) Encyclopedia of Cognitive Science, 4, pp. 524 – 527. London: Nature Publishing Group.

Intraub, H. (2004). Anticipatory spatial representation of 3D regions explored by sighted observers and a deaf-and-blind-observer. Cognition, 94(1), 19–37. 10.1016/j.cognition.2003.10.013

Intraub, H., Bender, R. S., & Mangels, J. A. (1992). Looking at pictures but remembering scenes.*Journal of Experimental Psychology: Learning*, Memory, and Cognition, 18(1), 180–191. 10.1037/0278-7393.18.1.180

Intraub, H., & Bodamer, J. L. (1993). Boundary extension: Fundamental aspect of pictorial representation or encoding artifact? *Journal of Experimental Psychology: Learning*, Memory, and Cognition, 19(6), 1387–1397. 10.1037/0278-7393.19.6.1387

Intraub, H., & Dickinson, C. A. (2008). False memory 1/20th of a second later: What the early onset of boundary extension reveals about perception. Psychological Science, 19(10), 1007–1014. 10.1111/j.1467-9280.2008.02192.x

Intraub, H., Gottesman, C. V., & Bills, A. J. (1998). Effects of perceiving and imagining scenes on memory for pictures. *Journal of Experimental Psychology: Learning*, Memory, and Cognition, 24(1), 186–201. 10.1037/0278-7393.24.1.186

Intraub, H., Morelli, F., & Gagnier, K. M. (2015). Visual, haptic and bimodal scene perception: evidence for a unitary representation. Cognition, 138, 132–147. 10.1016/j.cognition.2015.01.010

Intraub, H., & Richardson, M. (1989). Wide-angle memories of close-up scenes. Journal of experimental psychology. Learning, memory, and cognition, 15(2), 179–187. 10.1037//0278-7393.15.2.179

Intraub, H., Hoffman, J. E., Wetherhold, C. J., & Stoehs, S. A. (2006). More than meets the eye: the effect of planned fixations on scene representation. Perception & psychophysics, 68(5), 759–769. 10.3758/bf03193699

Johnson, M. R., & Johnson, M. K. (2014). Decoding individual natural scene representations during perception and imagery. Frontiers in human neuroscience, 8, 59. 10.3389/fnhum.2014.00059

Keogh, R., & Pearson, J. (2011). Mental imagery and visual working memory. PloS one, 6(12), e29221. 10.1371/journal.pone.0029221

Li, S., Zeng, X., Shao, Z., & Yu, Q. (2023). Neural Representations in Visual and Parietal Cortex Differentiate between Imagined, Perceived, and Illusory Experiences. The Journal of neuroscience, 43(38), 6508–6524. 10.1523/JNEUROSCI.0592-23.2023

Maguire, E. A., Intraub, H., & Mullally, S. L. (2016). Scenes, spaces, and memory traces. The Neuroscientist, 22(5), 432–439. 10.1177/1073858415600389

Mamus, E., & Boduroglu, A. (2018). The role of context on boundary extension. Visual Cognition, 26(2), 115–130. 10.1080/13506285.2017.1399947

Mullally, S. L., Intraub, H., & Maguire, E. A. (2012). Attenuated boundary extension produces a paradoxical memory advantage in amnesic patients. Current Biology, 22(4), 261–268. 10.1016/j.cub.2012.01.001

Munger, M. P., & Multhaup, K. S. (2016). No imagination effect on boundary extension. Memory & Cognition, 44, 73–88. 10.3758/s13421-015-0541-3

Oosterhof, N. N., Connolly, A. C., & Haxby, J. V. (2016). CoSMoMVPA: Multi-Modal Multivariate Pattern Analysis of Neuroimaging Data in Matlab/GNU Octave. Frontiers in neuroinformatics, 10, 27. 10.3389/fninf.2016.00027

Palmiero, M., Cardi, V., & Belardinelli, M. O. (2011). The role of vividness of visual mental imagery on different dimensions of creativity. Creativity Research Journal, 23(4), 372–375. 10.1080/10400419.2011.621857

Park, S., Intraub, H., Yi, D.-J., Widders, D., & Chun, M. M. (2007). Beyond the edges of a view: Boundary extension in human scene-selective visual cortex. Neuron, 54(2), 335–342. 10.1016/j.neuron.2007.04.006

Prince, J. S., Charest, I., Kurzawski, J. W., Pyles, J. A., Tarr, M. J., & Kay, K. N. (2022). Improving the accuracy of single-trial fMRI response estimates using GLMsingle. eLife, 11, e77599. 10.7554/eLife.77599

Ragni, F., Tucciarelli, R., Andersson, P., & Lingnau, A. (2020). Decoding stimulus identity in occipital, parietal and inferotemporal cortices during visual mental imagery. Cortex, 127, 371–387. 10.1016/j.cortex.2020.02.020

Reddy, L., Tsuchiya, N., & Serre, T. (2010). Reading the mind’s eye: decoding category information during mental imagery. NeuroImage, 50(2), 818–825. 10.1016/j.neuroimage.2009.11.084

Smith, S. M., & Nichols, T. E. (2009). Threshold-free cluster enhancement: addressing problems of smoothing, threshold dependence and localisation in cluster inference. NeuroImage, 44(1), 83–98. 10.1016/j.neuroimage.2008.03.061

Sreenivasan, K. K., Gratton, C., Vytlacil, J., & D’Esposito, M. (2014). Evidence for working memory storage operations in perceptual cortex. Cognitive, affective & behavioral neuroscience, 14(1), 117–128. 10.3758/s13415-013-0246-7

Stokes, M., Thompson, R., Cusack, R., & Duncan, J. (2009). Top-down activation of shape-specific population codes in visual cortex during mental imagery. The Journal of neuroscience : the official journal of the Society for Neuroscience, 29(5), 1565–1572. 10.1523/JNEUROSCI.4657-08.2009

Vaziri-Pashkam, M., & Xu, Y. (2017). Goal-Directed Visual Processing Differentially Impacts Human Ventral and Dorsal Visual Representations. The Journal of Neuroscience, 37(36), 8767–8782. 10.1523/JNEUROSCI.3392-16.2017

Wandell, B. A., Dumoulin, S. O., & Brewer, A. A. (2007). Visual field maps in human cortex. Neuron, 56(2), 366–383. 10.1016/j.neuron.2007.10.012

Xu Y. (2018). The Posterior Parietal Cortex in Adaptive Visual Processing. Trends in neurosciences, 41(11), 806–822. 10.1016/j.tins.2018.07.012

